# Deep generative modeling and clustering of single cell Hi-C data

**DOI:** 10.1101/2022.07.19.500573

**Authors:** Qiao Liu, Wanwen Zeng, Wei Zhang, Sicheng Wang, Hongyang Chen, Rui Jiang, Mu Zhou, Shaoting Zhang

## Abstract

Deciphering 3D genome conformation is important for understanding gene regulation and cellular function at a spatial level. The recent advances of single cell Hi-C technologies have enabled the profiling of the 3D architecture of DNA within individual cell, which allows us to study the cell-to-cell variability of 3D chromatin organization. Computational approaches are in urgent need to comprehensively analyze the sparse and heterogeneous single cell Hi-C data. Here, we proposed scDEC-Hi-C, a new framework for single cell Hi-C analysis with deep generative neural networks. scDEC-Hi-C outperforms existing methods in terms of single cell Hi-C data clustering and imputation. Moreover, the generative power of scDEC-Hi-C could help unveil the heterogeneity of chromatin architecture across different cell types. We expect that scDEC-Hi-C could shed light on deepening our understanding of the complex mechanism underlying the formation of chromatin contacts. scDEC-Hi-C is freely available at https://github.com/kimmo1019/scDEC-Hi-C.

**Key points:**

- scDEC-Hi-C provides an end-to-end framework based on autoencoder and deep generative model to comprehensively analyze single cell Hi-C data, including low-dimensional embedding and clustering.
- Through a series of experiments including single cell Hi-C data clustering and structural difference identification, scDEC-Hi-C demonstrates suprioir performance over existing methods.
- In the downstream analysis of chromatin loops from single cell Hi-C data, scDEC-Hi-C is capable of significantly enhancing the ability for identifying single cell chromatin loops by data imputation.

## Introduction

The rapid development in single-cell technologies enables us to reliably measure the genomic, transcriptomic and epigenomic features of a particular cellular context at single-cell resolution [1–4]. These powerful technologies provide scientists with the opportunity to study the unique patterns of cell type specificity and gene regulation. One fundamental question regarding the abundant single cell data is how to distinguish different cell types in a heterogeneous cell population based on the measured molecular signatures. A variety of computational approaches have been developed to decipher the heterogeneity across cell types based on transcriptome, methylome, and chromatin accessibility [5–11].

The majority of the current single-cell assays, such as RNA sequencing (scRNA-seq) and transposase-accessible chromatin using sequencing (scATAC-seq), ignore the spatial information of the genome, such as 3D chromatin structure, which plays an important role in genome functions, including gene transcription and DNA replication [12–14]. The emerging single cell Hi-C technologies bridge this gap by measuring the 3D chromatin structures in individual cells, which have the potential to comprehensively reveal the diverse genome functions underlying the unique genome structure [15–19].

Several computational methods have been proposed for the single cell Hi-C data analysis. For example, scHiCluster [20] introduced a random walk-based strategy for data imputation and used PCA for embedding. HiCRep/MDS [21] used multi-dimensional scaling (MDS) for learning a low-dimensional embedding. Higashi [22] is a recent method that utilized hypergraph representation learning for single cell imputation and embedding. However, all these methods require an additional clustering approach (e.g., K-means) for identifying cell types. In addition, choosing the most appropriate clustering approach is sometimes difficult as it is hard for a single clustering approach to perform the best across different datasets. Moreover, modeling the generation process of ultra-sparse single cell Hi-C data could help us better understand the formulation of 3D chromatin conformation, which was ignored by most previous methods.

To overcome the above mentioned limitations, we developed scDEC-Hi-C, a comprehensive end-to-end unsupervised learning framework for single cell Hi-C data embedding, clustering, and generation by deep generative neural networks. Unlike existing methods that treat embedding and clustering as two separated tasks, our approach enables simultaneously learning the low-dimensional embeddings of single cell Hi-C data and clustering the single cell Hi-C data by neural network in an unsupervised manner. From systematical experiments, scDEC-Hi-C demonstrates superiority in various tasks, including clustering the cell types, data imputation for quality enhancement, as well as data generation given a desired cell type. To the best of our knowledge, scDEC-Hi-C is the first computational framework that integrates the data embedding and clustering intrinsically for the single cell Hi-C data analysis.

## Results

### Overview of scDEC-Hi-C

scDEC-Hi-C consists of two major computational modules, including a convolutional autoencoder module for chromosome-wise representation learning and a deep generative module for cell-wise representation learning and clustering (Fig 1). The autoencoder module aims at extracting the low-dimensional features for each chromosome within a cell. Then the chromosome-wise features are transformed to cell-wise features through a chromosome readout function. We chose global concatenation for the readout function as default. The cell-wise generative model is adopted from our previous work scDEC [23] where G and H networks aim at bidirectional transformation between the m-dimensional latent space and n-dimensional representer space. Note that the latent variables ***z*** follows a standard Gaussian distribution N(**0,I**) and ***c*** follows a category distribution Cat(*K, **w***), which is parameterized by the number of clusters *K* and the weight ***w***. G network takes ***z*** and ***c*** as inputs and D_x_ network was used for matching the distribution of cell-wise representation *x* and G network output 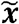 through adversarial training. Similarly, H network and D_z_ network also work in an adversarial manner where H network could learn the latent representation 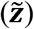 and infer the cluster 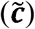 simultaneously. The detailed model architecture and training strategy can be found in Supplementary Table 1.

**Figure 1.**
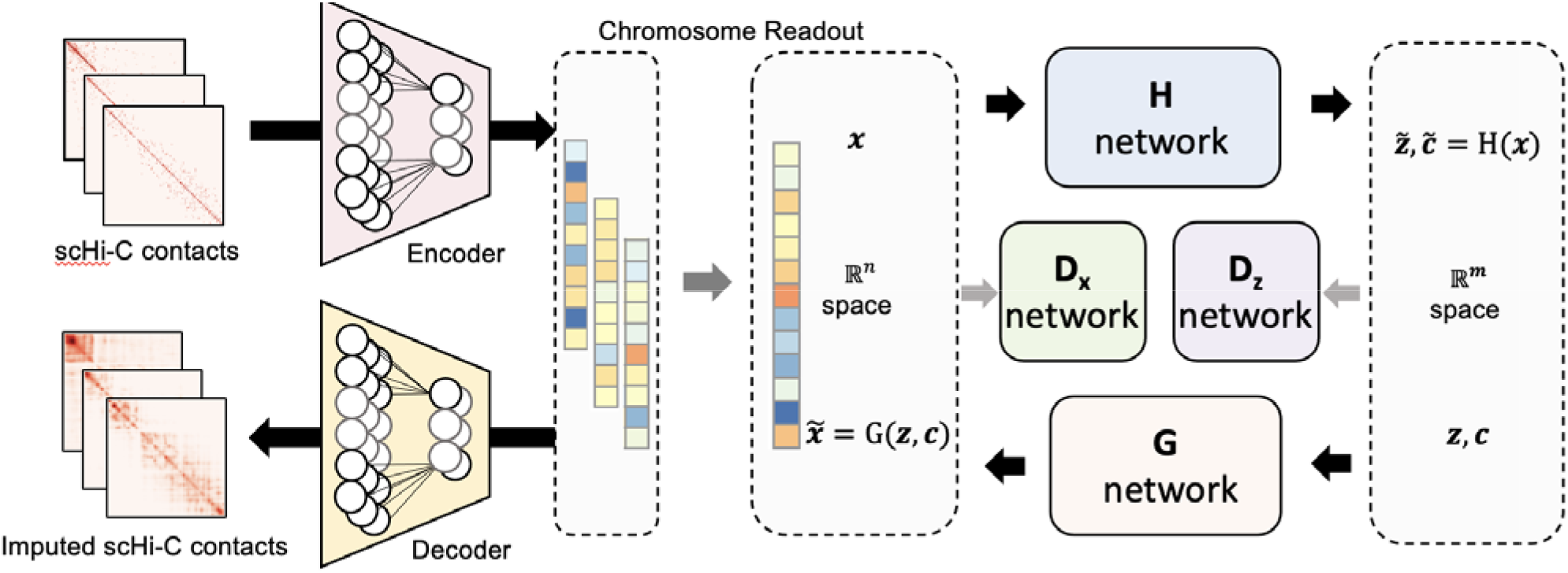
The overview of the proposed scDEC-Hi-C model. scDEC-Hi-C is a multi-scale model which contains a chromosome-wise convolutional autoencoder (CAE) and a cell-wise single cell deep embedding and clustering model. The intra-chromosome single-cell Hi-C contacts matrices are first fed to a CAE for dimension reduction and latent feature extraction. Then the chromosome readout (e.g., concatenation) is applied to get the cell-wise representation. The cell-wise deep generative neural networks can further learn a low dimensional representation of a cell and cluster each cell simultaneously. In the latent space, latent variables and sampled from a Gaussian distribution and a Category distribution respectively, are fed to the network. The network has two outputs of which one corresponds to the latent embedding ( ) and one corresponds to the estimated cluster label ( ). The and discriminator networks are used for adversarial training.

### scDEC-Hi-C is capable of identifying cell heterogeneity

A fundamental problem in single cell Hi-C data analysis is to identify different cell types in heterogeneous cell populations. To evaluate the performance of scDEC-Hi-C on this task, we adopted two commonly used benchmark datasets here and systematically compared scDEC-Hi-C to three baseline methods (see Methods for data preprocessing and Supplementary Table 2). Three metrics, including NMI, ARI, and Homogeneity, were introduced for measuring the performance in this unsupervised learning task in order to quantify the ability for distinguishing different cell types in the single cell Hi-C datasets (see Methods). Note that all baseline methods are only able to learn the embedding for each single cell and require additional clustering methods (e.g, K-means) while scDEC-Hi-C simultaneously learns cell embeddings and assigns clustering labels to each cell. scDEC-Hi-C is capable of learning embeddings which could separate cells from different cell types with a relatively larger margin than other baseline methods (Fig.2A-B). It is worth mentioning that scDEC-Hi-C exhibits superior performance on Ramani dataset [17] by outperforming other methods with an ARI of 0.845, compared to 0.826 of Higashi, 0.795 of scHiCluster, and 0.785 of HiCRep/MDS (Fig. 2C). In the second Dip-C dataset [24] where only annotated labels were available, we treat the annotated labels as surrogate ground truth labels. All methods demonstrate significantly lower clustering performance than the Ramani dataset with ground truth label. Specifically, scDEC-Hi-C demonstrates slightly lower performance than Higashi (Fig. 2C). In the readout module in scDEC-Hi-C, the information coming from each chromosome was aggregated. Thus it is worthy to evaluate the contribution of each chromosome. The experimental results show that chromosome 11 contributed the most in Ramani dataset and scDEC-Hi-C consistently outperformed Higashi in 18 chromosomes out of 24 (Fig. 2D). To further investigate the effect of sequencing depth on the clustering performance, we randomly dropout the sequencing reads with different rate. scDEC-Hi-C consistently outperforms all other baseline methods at different dropout rates (Fig. 2E).

**Figure2.**
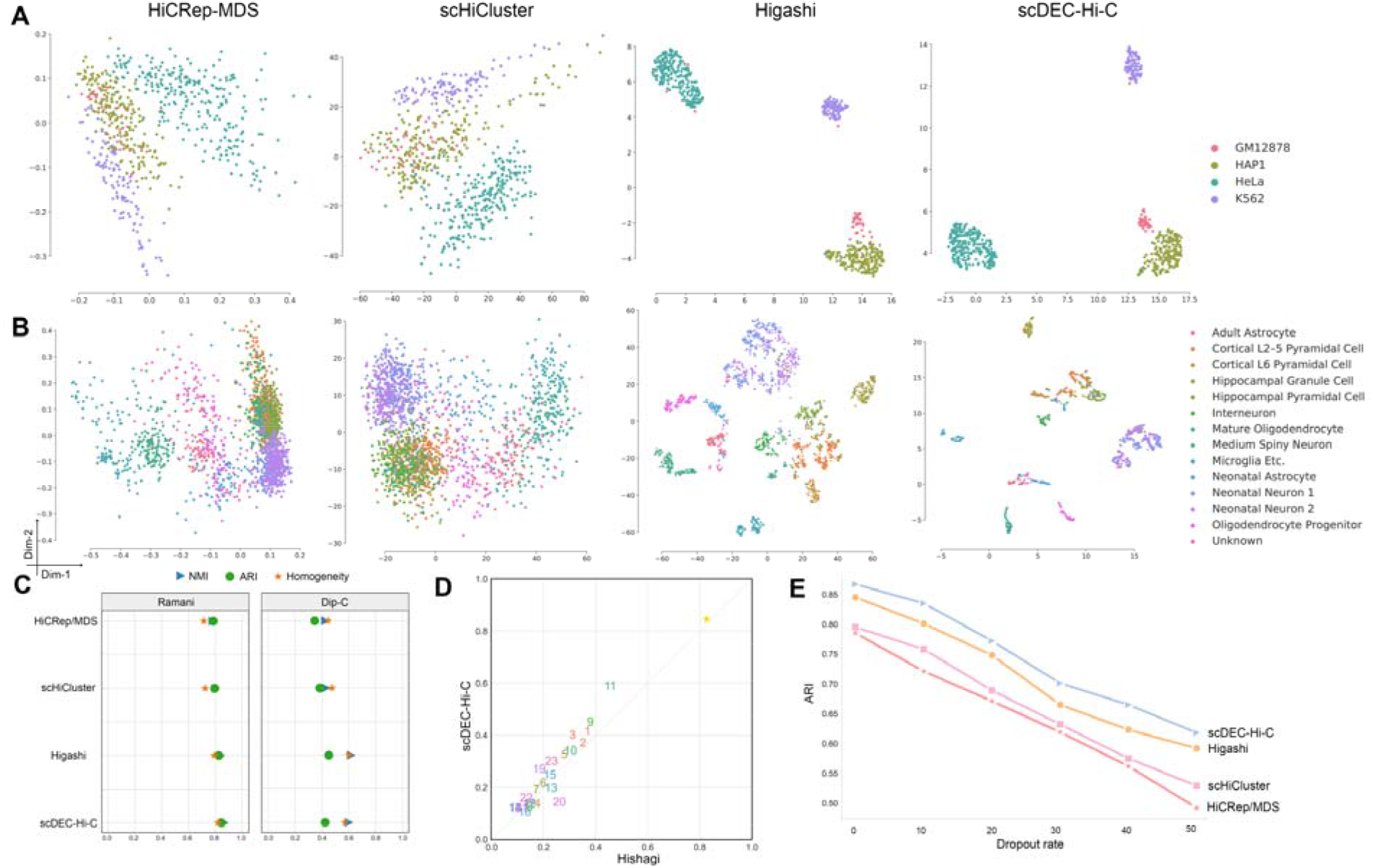
The performance of scDEC-Hi-C method and baseline methods on single cell Hi-C datasets. (A) The embeddings visualization of Ramani dataset across four methods. (B) The embeddings visualization of Dip-C dataset across four methods. (C) The clustering performance in terms of NMI, ARI and Homogeneity of four methods across two datasets. (D) The performance of scDEC-Hi-C and baseline methods under different dropout rate on Ramani dataset.

### scDEC-Hi-C enables the identification of structural differences

In single cell Hi-C data analysis, one fundamental question to ask is whether cell type specificity is revealed by the structural difference regions in Hi-C contacts. The cell type specificity in single cell data, such as single cell RNA-seq and single cell ATAC-seq, can be clearly revealed by marker genes or differential peaks [25]. In bulk Hi-C data, it has been validated that cell type specificity is highly associated with the dynamic chromatin loops within topologically associating domains (TADs) [26, 27]. Therefore, it is worthwhile to investigate whether the structural differences also exist in single cell Hi-C data. To explore this, we used the autoencoder from the first stage of scDEC-Hi-C model as an approach for scDEC-Hi-C imputation. In brief, we segmented Hi-C contact matrix of each chromosome per cell into non-overlapping square patches within the range of 1Mbp. We then treated the output of decoder as the imputed single cell Hi-C data (see Methods). We designed extensive experiments to evaluate whether the imputed single cell Hi-C data could reveal more biological insights than the raw data. We aggregated single cells of K562 and GM12878 cell lines from Ramani dataset and then merged them as the aggregated Hi-C data. In the meanwhile, we also downloaded the bulk Hi-C data from GM12878 and K562 cell lines as ground truth for validation. From the Hi-C profile of a genomic region (chr9: 132.9M-134.9M), K562 and GM12878 have significantly different Hi-C contacts map the difference is also emphasized by the imputed single cell Hi-C data (Fig. 3A). Specifically, the chromatin structural boundaries marked by the rectangle is much clearer by imputed data than the raw data, which demonstrates the power and effectiveness of scDEC-Hi-C in enhancing the resolution of chromatin structural boundaries. It is also noticeable that the chromatin structural boundaries revealed by bulk Hi-C data have a larger consistency with imputed single cell data than the raw single cell data. To further investigate the regulatory landscape of this genomic region, we downloaded both RNA-seq and histone modification data from ENCODE database [28] and visualized them with the help of WashU Epigenome Browser [29]. It can be seen that both RNA-seq signal and H3K4me1 marker are more enriched in K562 cell line than GM12878 cell line in the bounded region (Fig. 3B), which indicates a strong activity of regulatory elements such as enhancer in K562. Next, we designed quantitative experiments to verify whether the resolution of single Hi-C data could be improved by scDEC-Hi-C model. Taking the bulk K562 Hi-C data as ground truth, we calculated the Pearson’s correlation of Hi-C interactions of different distances between ground truth and imputed data. It is seen that the interactions at a larger distance are more difficult to impute (Fig. 3C). The correlation between bulk Hi-C data and raw single cell Hi-C data is less than 0.25 while the single cell Hi-C data imputed by scDEC-Hi-C and scHiCluster are much higher than the baseline. scDEC-Hi-C consistently outperforms scHiCluster at different distances ranging from 0 to 1Mb. To sum up, scDEC-Hi-C enables improving the identification of structural boundaries which further helps us study the chromatin structure difference across diverse cell types.

**Figure3.**
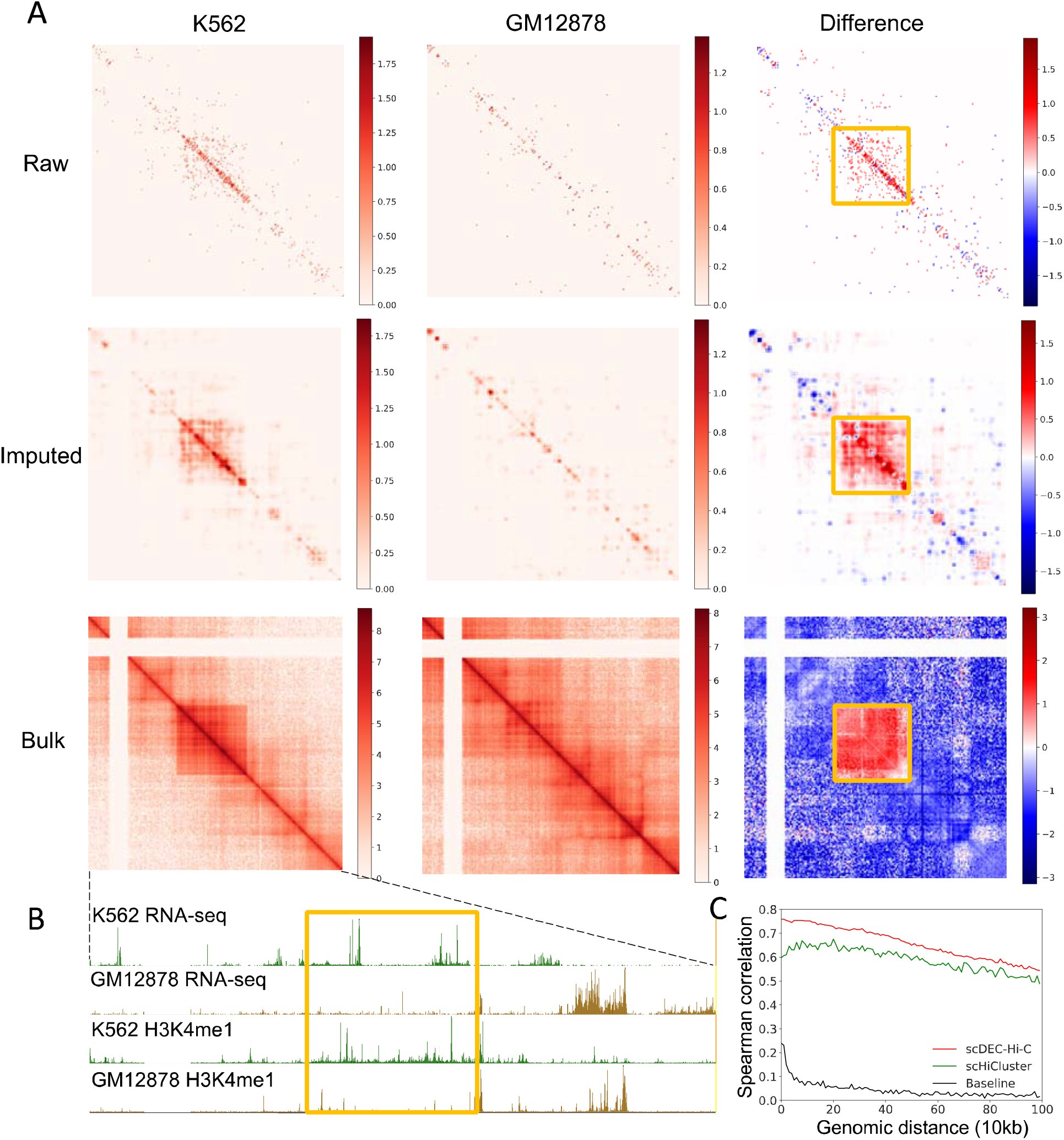
The imputation results of scDEC-Hi-C method. (A) The first row denotes merged single cell Hi-C profile of 40 cells of a genomic region (chr9: 132.9M-134.9M) across two diverse cell lines. The middle row denotes the corresponding imputed single cell Hi-C profile with scDEC-Hi-C. The third row denotes the corresponding bulk Hi-C profile of the two cell lines. The differences of the Hi-C profile from two cell lines are illustrated. (B) Genome annotation including RNA-seq and H3K4me1 histone marker across two cell lines of the same genomic region. (C) The Spearman correlation between bulk K562 Hi-C data and aggregated single cell Hi-C data after imputation by scDEC-Hi-C (red) and scHiCluster (green). The baseline (black) denotes the Spearman correlation between bulk K562 Hi-C data and aggregated single cell Hi-C data without imputation.

### scDEC-Hi-C enhances the discovery of chromatin loops

Chromatin loops are defined as a pair of genomic regions that are brought into spatial proximity, which can be inferred from bulk Hi-C data. Chromatin loops have been proved to be highly relevant to gene regulation, cell fates and functions. We then intended to explore whether the chromatin loops can also be identified within single cell Hi-C data. Similarly, we merged single cell Hi-C data of K562 and GM12878 cell lines, respectively. In the meanwhile, we also downloaded the corresponding bulk Hi-C data for comparison. We applied Fit-Hi-C [30], a computational tool for calling chromatin loops from Hi-C data, to bulk Hi-C data and imputed single cell Hi-C data by scDEC-Hi-C, respectively. There are 6478 chromatin loops in GM12878 cell line while 732 (11.3%) chromatin loops are also discovered in imputed single cell Hi-C data (Fig. 4A). scDEC-Hi-C additionally identified 294 chromatin loops which are not contained in the bulk Hi-C chromatin loops. Note that only 196 chromatin loops can be identified from raw single cell Hi-C data and scDEC-Hi-C significantly improves the precision from 1.4% to 11.3% by imputation (Supplementary Figure 1). We visualized the chromatin loops in a genomic region (chr3:118.2M-120.2M) of bulk Hi-C chromatin loops versus either raw single cell Hi-C data (Fig. 4B) or imputed single cell Hi-C data (Fig. 4C). In K562 cell line, 12.0% of the chromatin loops from bulk Hi-C data can be also recovered by imputed single cell Hi-C data and 72.5% of the chromatin loops from imputed single cell Hi-C data are also contained in bulk chromatin loops (Fig. 4D). In the same genomic region, imputed single cell Hi-C data contains three chromatin loops while two of them were consistent with bulk Hi-C chromatin loops (Fig. 4F) while the raw single cell Hi-C data only has one false chromatin loop (Fig. 4E). To conclude, scDEC-Hi-C is able to promote the identification of chromatin loops from Hi-C data.

**Figure 4.**
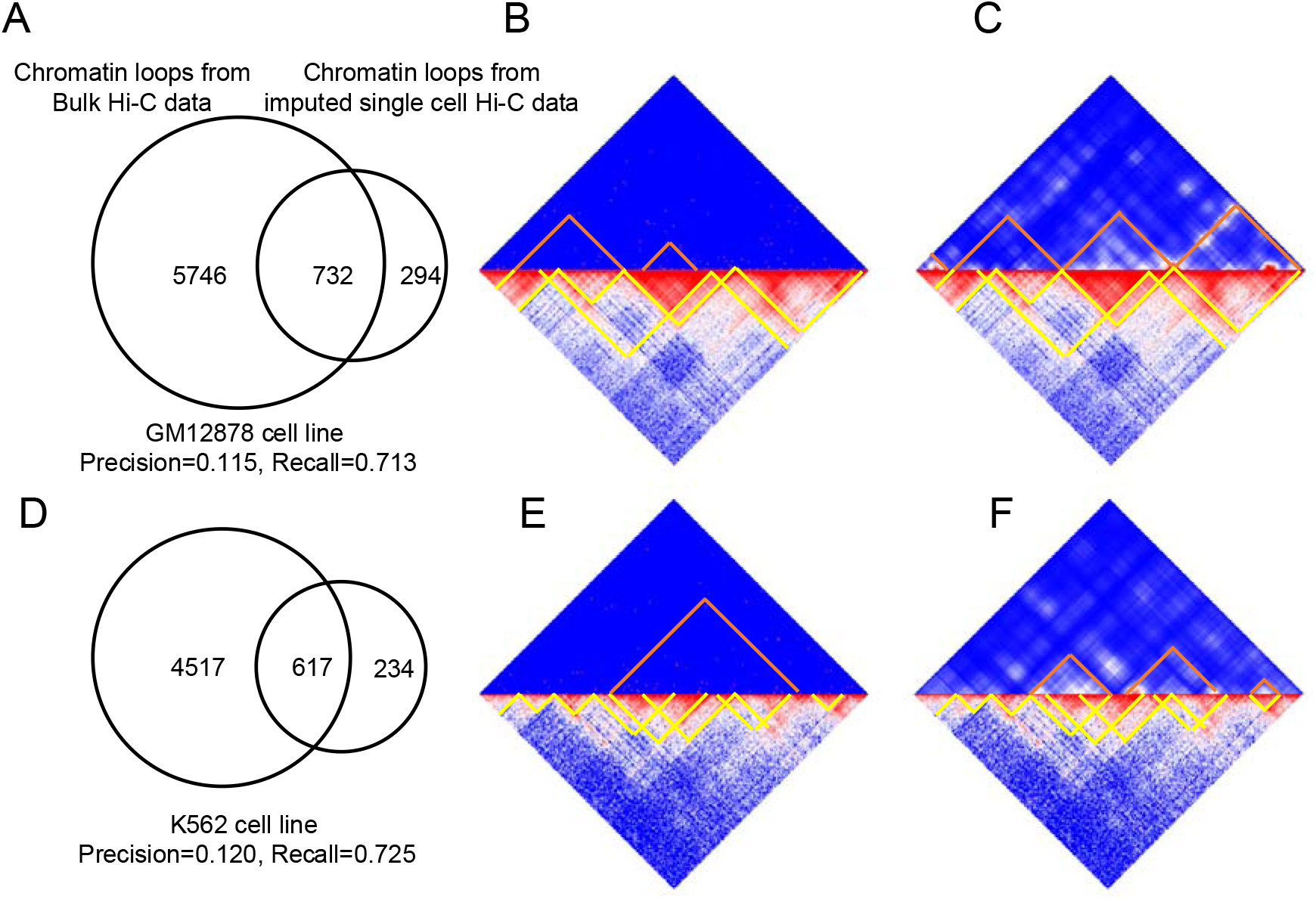
scDEC-Hi-C facilitates the identification of chromatin loops. (A) The Venn plot of chromatin loops from bulk Hi-C data and single-cell Hi-C data imputed by scDEC-Hi-C in GM12878 cell line. (B) The chromatin loops from raw single cell Hi-C data versus chromatin loops from bulk Hi-C data of a GM12878 cell line genomic region (chr3:118.2M-120.2M). (C) The chromatin loops from imputed single cell Hi-C data versus chromatin loops from bulk Hi-C data in GM12878 cell line of the same genomic region. (D) The Venn plot of chromatin loops from bulk Hi-C data and single-cell Hi-C data imputed by scDEC-Hi-C in K562 cell line. (E) The chromatin loops from raw single cell Hi-C data versus chromatin loops from bulk Hi-C data of a K562 cell line genomic region (chr3:118.2M-120.2M). (F) The chromatin loops from imputed single cell Hi-C data versus chromatin loops from bulk Hi-C data in in GM12878 cell line of the same genomic region.

### Ablation analysis

To systematically evaluate the robustness of scDEC-Hi-C, we designed the following ablation studies. We used Ramani dataset for the ablation studies. First, we removed the cell-wise scDEC module and only kept the chromosome-wise convolutional autoencoder module. We directly used K-means for clustering the features from concatenated autoencoder features. The ARI, NMI and Homogeneity decreases by 6.2%, 7.2%, and 7.1%, respectively. Second, we trained the chromosome-wise autoencoder model first and then fixed the weights in the autoencoder and trained the cell-wise scDEC module. Without joint training of the multi-stage modules, the performance also decreases by 2.4% of ARI, 2.8% of NMI and 2.5% of Homogeneity. The model ablation studies demonstrate the significant contribution of both multi-stage model and joint training strategy.

**Table 1.** Model ablation studies. The standard deviation of the metric was calculated based on five runs.

|  | ARI | NMI | Homogeneity |
| --- | --- | --- | --- |
| scDEC-Hi-C | 0.845±0.010 | 0.867±0.012 | 0.819±0.009 |
| scDEC-Hi-C w/o scDEC | 0.783±0.009 | 0.795±0.015 | 0.748±0.014 |
| scDEC-Hi-C w/o joint training | 0.821±0.013 | 0.839±0.017 | 0.794±0.013 |

### Conclusion and discussion

In this study, we proposed scDEC-Hi-C, a computational tool for comprehensive single cell Hi-C data analysis using deep generative neural network. Unlike previous works that treat dimension reduction and clustering of the single cell Hi-C data as two separated and independent tasks, scDEC-Hi-C intrinsically integrates the task of learning a low-dimensional representation and clustering the single cell by designing a two-stage multi-scale framework, which is composed of a chromosome-wise autoencoder and a cell-wise symmetric GAN model. During the training, the multi-scale models are simultaneously optimized and the results of embedding and clustering are benefitting each other. Based on a series of experiments, scDEC-Hi-C achieves superior or competitive performance compared to state-of-the-art baseline methods. For the downstream analysis, scDEC-Hi-C model demonstrated the excellent ability of imputing the sparse and noisy single cell Hi-C data, which facilitates the identification of chromatin structural differences and chromatin loops. Besides, scDEC-Hi-C also shows the superior power in generating the Hi-C profile of different cell types, which has been confirmed to be consistently with the cell type label (Supplementary Figure 2).

We also provide several directions for further improving our work. First, the inter-chromosomal interactions, which were ignored by existing methods and scDEC-Hi-C, have been proved to regulate gene expression [31]. Second, incorporating multi-omics data, including functional genomic regulatory annotation data [32, 33] and pharmaceutical interaction data [34, 35], could potentially improve the performance. Third, it is worthwhile for applying scDEC-Hi-C to other different types of 3D genome interaction data such as HiChIP [36].

With scDEC-Hi-C, researchers can perform single cell Hi-C experiments of the cell types or tissues with interest. Then one can simultaneously perform unsupervised learning analysis on single cell Hi-C data and uncover biological findings through the imputation and generation power. We expect that scDEC-Hi-C can help unveil the single cell regulation mechanism in 3D genome.

## Methods

### Data preprocessing

For Ramani dataset, we filtered cells with less than 5000 contacts. Then we collected 624 cells for Ramani dataset. For Dip-C dataset, we used the same QC strategy from the original paper[24] and collected 1954 annotated cells across 14 cell types. The details of datasets were summarized in Supplementary Table 2. The raw Hi-C contact matrices were log-transformed and then resized by spine interpolation so that the Hi-C contact matrix of each chromosome was represented as a 50 by 50 matrix. Then we applied a mean filtering and random walk as suggested by scHiCluster [20]. The chromosome-wise module encodes each chromosome into a 50-dimensional vector and then concatenated across all chromosomes. The cell-wise module further learns a low-dimensional representation of a cell with dimension of latent variable ***z*** set to 10. The embedding of each cell was based on the concatenation of reconstructed 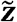 and 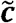 (before softmax).

### Adversarial training in scDEC-Hi-C model

The scDEC-Hi-C is multi-scale unsupervised learning model derived from our previous works Roundtrip and scDEC [23, 37] with extensive modifications. scDEC-Hi-C mainly contains a chromosome-wise module convolutional autoencoder (CAE) [38] and a cell-wise model scDEC. The CAE module aims at mapping scHi-C data from the original data space to a representer space, which significantly reduced the data dimension. Specifically, the CAE module takes the single cell Hi-C interaction of each chromosome as a training instance and each intra-chromosomal interaction matrix will be encoded to a fixed dimension vector through encoder. The embedding vectors for intra-chromosomal interaction matrices within each cell are concatenated in the representer space to obtain a fused embedding. The training of the CAE can be formulated as

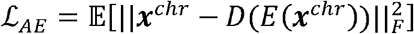

where *x^chr^* denotes an intra-chromosomal Hi-C interaction matrix and *E*(·), D(·) denote the encoder and decoder in the CAE module, respectively. The chromosome-wise features E(*x^chr^*) of each chromosome were concatenated to obtain the cell-wise representation by

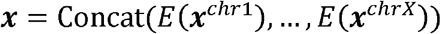

The scDEC module takes cell-wise fused embedding in the representer space as input and learns the low-dimension embedding of a cell in the latent space and clusters the cells simultaneously. scDEC module is composed of a pair of two GAN models. For the forward GAN model, a pair of latent variables ***z*** and ***c*** are sampled from a Gaussian distribution and a Categorical distribution, respectively. The categorical distribution is updated through an adaptive mechanism (Supplementary Table 3). G network is used for conditionally generating fake data 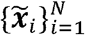 that have a similar distribution to the real data 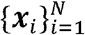 in the representer space while the discriminator network D_x_ tries to discern true data from generated samples in the representer space. In the backward GAN model, the function H and the discriminator D_z_ aim at transforming the data from representer space to the latent space. Discriminators can be considered as binary classifiers where any input data point will be asserted to be positive or negative. Besides, we used WGAN-GP [39] as the architecture for the pair of GAN models where the gradient penalties of discriminators were considered as additional loss terms. We then define the objective loss functions of the above four networks (G, H, D_x_ and D_z_) in the training process as

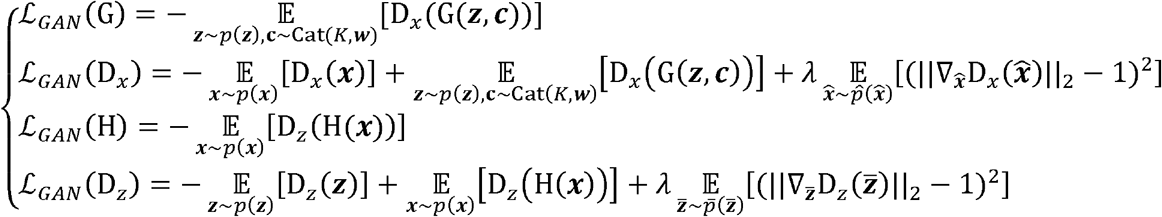

where p(***z***) and Cat(*K, **w***) denote the distribution of the continuous variable and discrete variable in the latent space. In practice, sampling ***x*** from *p*(***x***) can be regarded as a process of randomly sampling from *i.i.d* data in the representer space with replacement. 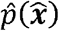 and 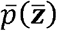 denote a uniformly sampling from the straight line between a pair of points sampled from true data and generated data in the representer and latent space, respectively. *λ* is a penalty coefficient which is set to 10 in all experiments.

### Roundtrip loss

During the training process, we also intend to minimize the roundtrip loss [37] which is defined as ρ((***z, c***), H(G(***z,c***))))and ρ(***x***, G(H(***x***))) where ***z*** and ***c*** are sampled from p(***z***) and Cat(*K, **w***), respectively. The basic principle for this loss is to minimize the distance when a data point goes through a roundtrip transformation between two different data domains. Specifically, we applied *l_2_*loss to the continuous part in roundtrip loss and cross entropy loss to the discrete part in roundtrip loss. We further denoted the roundtrip loss as

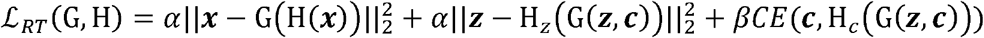

where *α* and *β* are two coefficients and are both set to 10 in the experiments. H_z_(·) and H_c_(·) denote the continuous and discrete part of H(·), respectively. *CE*(·) represents the cross-entropy function. The idea of roundtrip loss which exploits transitivity for regularizing structured data has also been used in previous works [40, 41].

### Joint training

Combining the adversarial training loss and roundtrip loss together, we can get the full training loss for the scDEC module as 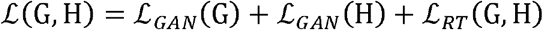 and 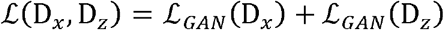, respectively. We iteratively updated the weight parameters in two generative models (G and H) and the two discriminative models (D_x_ and D_z_), respectively. Thus, the training of scDEC module can be represented as

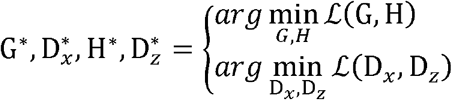

To further achieve joint training of CAE and scDEC modules, we first pretrained the CAE module for 100 epochs. Then we updated the parameters of CAE and scDEC iteratively. The Adam optimizer [42] with a learning rate of 2 × 10^-4^ was used for optimizing the parameters in neural networks. The whole training process is illustrated in Supplementary Table 4 in detail.

### Data imputation by scDEC-Hi-C model

We use the chromosome-wise model autoencoder for data imputation. Specifically, the reconstructed Hi-C map from the decoder was regarded as the imputed single cell Hi-C data. We used the same strategy in [13] for Hi-C matrices extraction.

### Data generation by scDEC-Hi-C model

We generate the intermediate cell state (embeddings) of single cell Hi-C data by interpolating the latent indicator *c* of two “neighboring” cell types. Assume that two cell types correspond to the latent indicator ***C*_1_** and ***c*_2_**, respectively. The generated single cell Hi-C profile can be represented as G(***z,ĉ***) where **ĉ** = α***C_1_*** + (1 — α)***c_2_***.Note that the *α* is the coefficient from 0 to 1 and ***z*** is sampled from a standard Gaussian distribution.

### Network architecture in scDEC-Hi-C

For the CAE module, the encoder contains four convolutional layers and two fully connected layers while the decoder consists of two fully connected layers and four transposed convolutional layers for reconstructing the Hi-C interaction matrices. For the scDEC module. The G network contains ten fully connected layers and each hidden layer has 512 nodes while the H network contains ten fully-connected layers and each hidden layer has 256 nodes. D_x_ and D_z_ both contain two fully connected layers and 256 nodes in the hidden layer. Note that batch normalization [43] was used in discriminator networks.

### Updating the Category distribution

The probability parameter ***w*** in the Category distribution Cat(*K, **w***) is adaptively updated every 200 batches of data based on the inferred cluster label (Supplementary Table 3).

### Evaluation metrics for clustering

We compared different methods for clustering according to three commonly used metrics, normalized mutual information (NMI) [44], adjusted Rand index (ARI) [45] and Homogeneity [46]. Assuming that *U* and *V* are true label assignment and predicted label assignment given *n*observation data points, which have *C_U_* and *C_V_* clusters in total, respectively. NMI is then calculated as

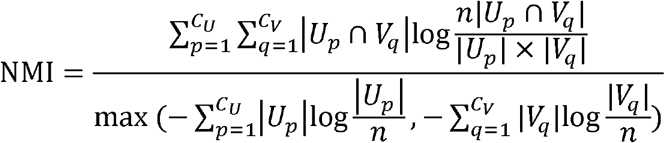

The Rand index [47] is a measure of agreement between two cluster assignments while ARI corrects lacking a constant value when the cluster assignments are selected randomly. We define the following four quantities: 1) *n_1_*: number of pairs of two objects in the same groups in both *U*and *V*, 2) n_2_: number of pairs of two objects in different groups in both *U* and *V*, 3) n_3_: number of pairs of two objects in the same group of *U* but different group in *V*, 4) n_4_: number of pairs of two objects in the same group of *V* but different group in U. Then ARI is calculated by

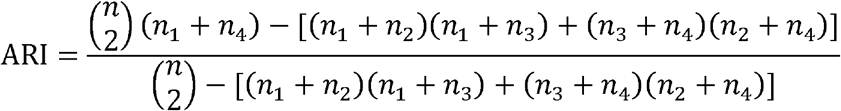

Homogeneity is calculated by 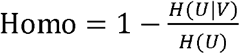, where

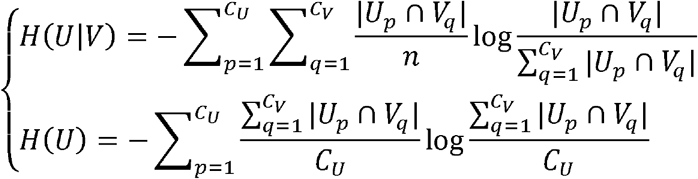

### Baseline methods

We compared scDEC-Hi-C to three comparison methods in our study. scHiCluster is a PCA-based method that could be used for imputing and clustering scHi-C data. scHiCluster was implemented from https://github.com/zhoujt1994/scHiCluster and the default parameters were used. HiCRep/MDS used multidimensional scaling to embed scHi-C data into two dimension and was implemented from https://github.com/liu-bioinfo-lab/scHiCTools. Higashi is a hypergraph representation learning framework for embedding scHi-C data. We downloaded Higashi from https://github.com/ma-compbio/Higashi and implemented using the default parameters.

## Data availability

Three datasets were used in this study. scHi-C dataset of four human cell lines (GM12878, HAP1, HeLa and K562) was collected from Ramani et al (GEO: GSE84920). scHi-C dataset of mouse brain development was collected from Tan et al (GEO: GSE162511). Note that the first dataset has ground truth cluster label for each cell. The latter dataset only contains annotated labels, which were used as surrogate labels in the clustering experiments.

## Code availability

scDEC-Hi-C is an open-source software based on the TensorFlow library [48], which can be downloaded from https://github.com/kimmo1019/scDEC-Hi-C.

## Acknowledgement

The authors thank the anonymous reviewers for their valuable and constructive suggestions.

## Author contributions statement

Q.L., R.J., M.Z. and S.T.Z. conceived the study. Q.L. designed and implemented scDEC-Hi-C. Q.L. and W.W.Z. performed data analysis. Q.L. and W.W.Z. interpreted the results. Q.L. wrote the manuscript. Other authors provided editorial support. All authors read and approved the final version of the manuscript.

