## Supplementary Data for "Deep generative modeling and clustering of single cell Hi-C data"

**Supplementary Figures**

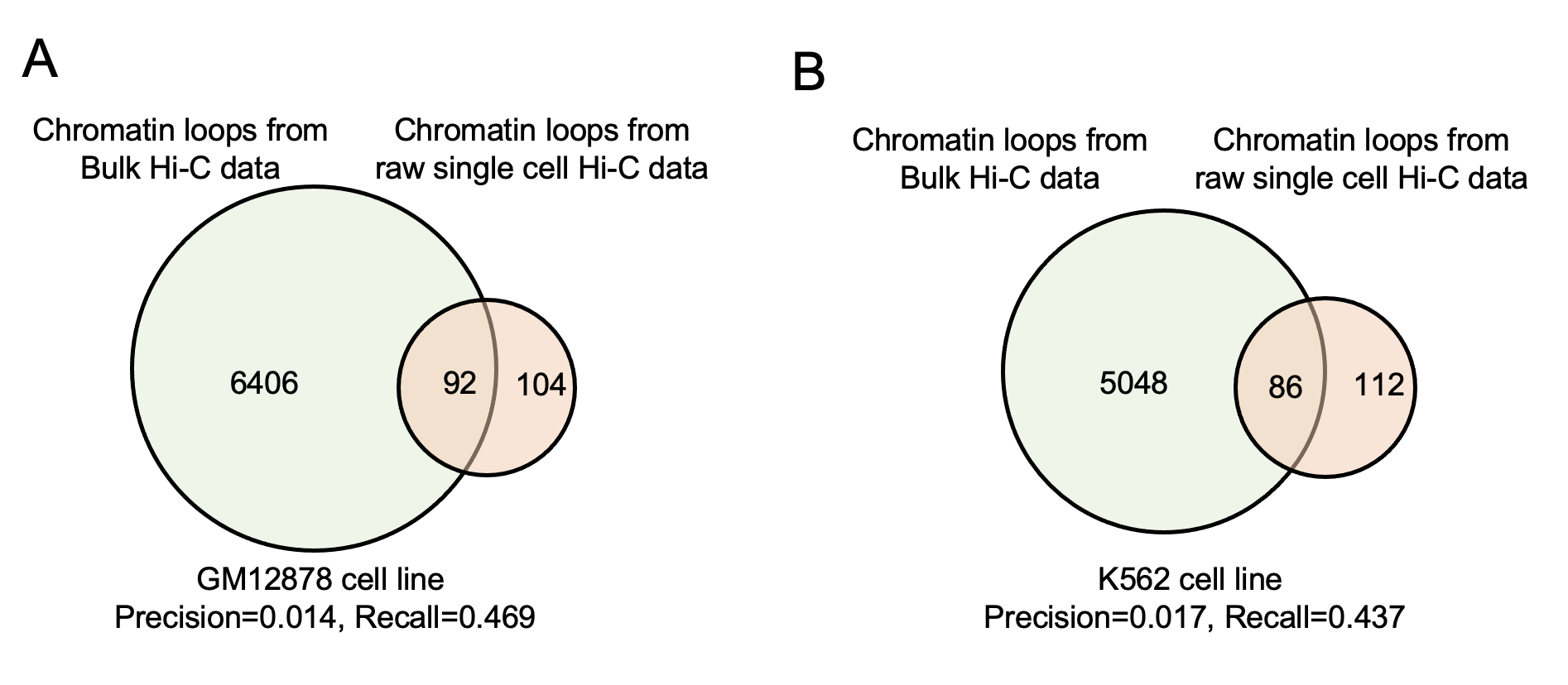

**Supplementary Figure 1.** The chromatin loops called from bulk Hi-C data versus chromatin loops called from raw single cell Hi-C data from GM12878 cell line and K562 cell line, respectively.

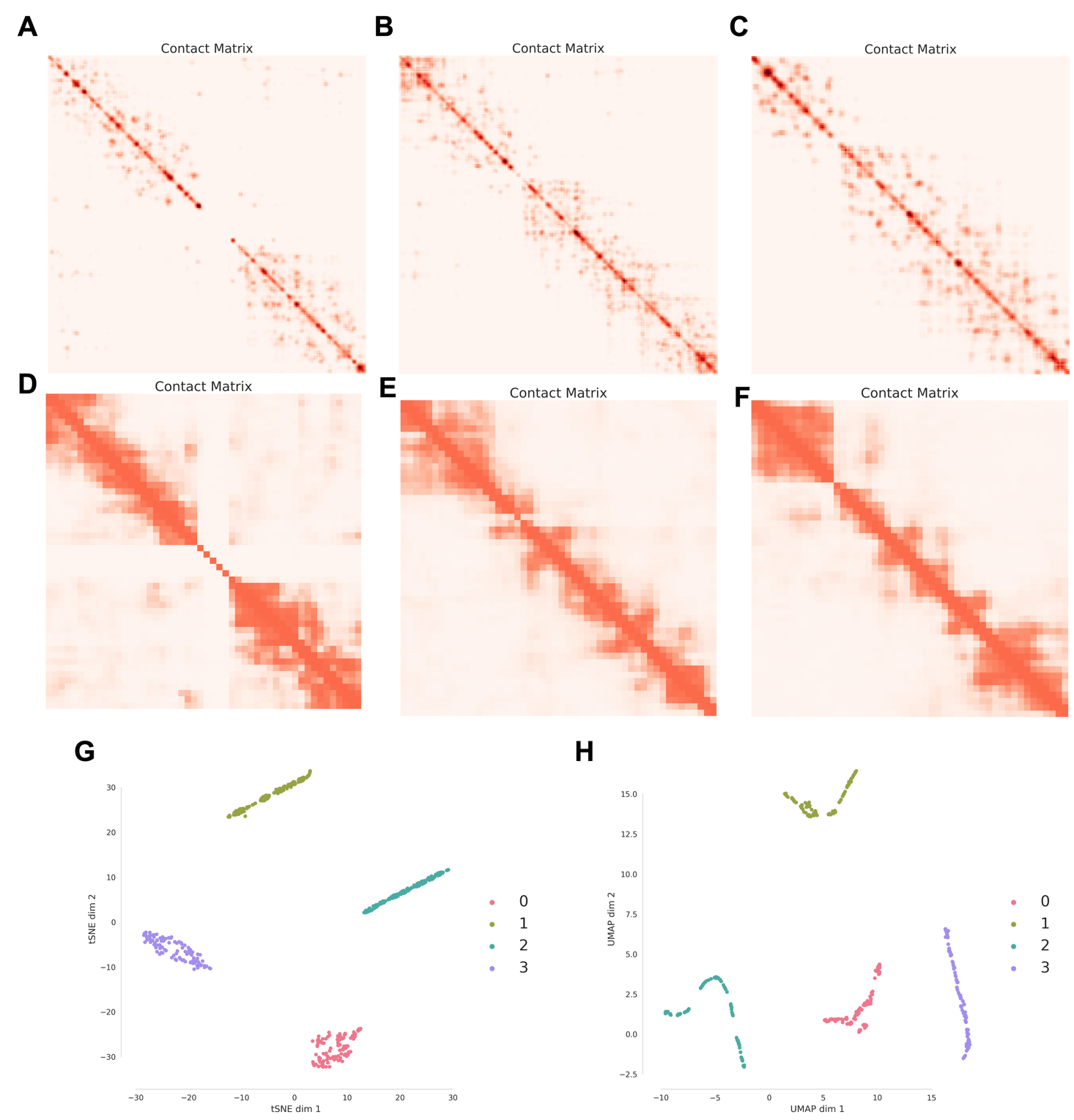

**Supplementary Figure 2.** The single cell Hi-C maps generated by scDEC-Hi-C. (A-C) True single cell Hi-C data of Chr1, Chr2, and Chr5 from the average of GM12878 cells. (D-F) Generated single cell Hi-C data on Chr1, Chr2, Chr5 of GM12878 cell line. (G-H) The t-SNE and UMAP visualization of the G network output by setting the latent variable c to four different categories, respectively (each category contains 100 points, denoting 100 generated cell profile). Label ‘0’, ‘1’, ‘2’, and ‘3’ represents 'GM12878', 'HAP1', 'HeLa', 'K562', respectively.

**Supplementary Tables**

**Supplementary Table 1**. The detailed hyperparameters of scDEC-Hi-C model. FC: fully-connected layer, ReLu: rectified linear unit, Identity: identity mapping (without non-linear activation function), Tanh: Tangent Hyperbolic function, Softmax: softmax function, batchnorm: batch normalization. Conv: convolution, Deconv: deconvolution (also known as transposed convolution). The hyperparameter optimization was done by grid searching of several key hyperparameters, including the learning rate (1e-3, 1e-4, 2e-4,1e-5), number of layers (2, 5,10,15) and number of hidden nodes (128, 256, 512) in each network (G, H, D_z_ and D_x_).

| **Encoder** E | **Decoder** D |
| --- | --- |
| Input $\boldsymbol{x}^{chr}\in\mathbb{R}^{112\times112}$ | Input $\boldsymbol{z}^{chr}\in\mathbb{R}^{50}$ |
| (Conv, MaxPooling, batchnorm)$\times4$ | Reshape |
| Flatten, FC | (Deconv, batchnorm)$\times4$ |
| Output $\boldsymbol{z}^{chr}\in\mathbb{R}^{50}$ | Output ${\hat{\boldsymbol{x}}}^{chr}\in\mathbb{R}^{112\times112}$ |
| **Generator** G | **Discriminator** D*_x_* |
| Inputs $\boldsymbol{z}\in\mathbb{R}^{d}$ and $\boldsymbol{c}\in\mathbb{R}^{K}$ | Input $\boldsymbol{x}\in\mathbb{R}^{20}$ or $\tilde{\boldsymbol{x}}\in\mathbb{R}^{20}$ |
| Concat($\boldsymbol{z}$**,**$\boldsymbol{c}$) | FC, 256, ReLu |
| (FC, 512, ReLu)$\times10$ | FC, 256, batchnorm, Tanh |
| FC, 20, Identity | Fc, 1, Identity |
| Output $\tilde{\boldsymbol{x}}\in\mathbb{R}^{20}$ | Output ${logit}_{x}\in\mathbb{R}^{1}$ |
| **Generator** H | **Discriminator** D*_z_* |
| Input $\boldsymbol{x}\in\mathbb{R}^{50*nb\_chrom}$ | Input $\boldsymbol{z}\in\mathbb{R}^{d}$ or $\tilde{\boldsymbol{z}}\in\mathbb{R}^{d}$ |
| (FC, 256, ReLu)$\times10$ | FC, 256, ReLu |
| FC, *d*, Identity and FC, *K*, Softmax | FC, 256, batchnorm, Tanh |
| Outputs $\tilde{\boldsymbol{z}}\in\mathbb{R}^{d}$ and $\tilde{\boldsymbol{c}}\in\mathbb{R}^{K}$ | Fc, 1, Identity |
|  | Output ${logit}_{z}\in\mathbb{R}^{1}$ |

**Supplementary Table 2**. The detailed of the single cell/bulk Hi-C datasets used in this study.

| Dataset | No. Of cells | Median reads | No. of cell types | Accession |
| --- | --- | --- | --- | --- |
| Ramani | 624 | 18529.5 | 4 | GSE84920 |
| Dip-C | 1954 | 312299 | 14 | GSE162511 |

| Dataset | File name | No. of original loops | Accession |
| --- | --- | --- | --- |
| Rao-GM12878 | GSE63525_GM12878_primary_intrachromosomal_contact_matrices.tar.gz | 8055 | GSE63525 |
| Rao-K562 | GSE63525_K562_intrachromosomal_contact_matrices.tar.gz | 6058 | GSE63525 |

**Supplementary Table 3.** Adaptively updating $w$ in the Category distribution in the latent space of sDEC-Hi-C model. $r$ is the ratio coefficient and $\epsilon$ denotes the lower bound of the cluster proportion. $U\left( p,q \right)$ represents a uniform distribution between $p$ and $q$.

**Algorithm** Adaptively updating $w$

**Input**: $\boldsymbol{w}^{(t)}$ and $\{{\tilde{\boldsymbol{c}}}_{i}|i=1,\ldots,N\}$ $r=0.2$, $\epsilon=0.02;$

**Output**:$\boldsymbol{w}^{(t+1)}$;

**For** $k\leftarrow1$ to $K$ **do**

$\boldsymbol{w}^{esk}\left( k \right)=\sum_{i=1}^{N} I(\mathrm{argmax}\left( {\tilde{\boldsymbol{c}}}_{i} \right)=k)/N;$

**end**

$$\boldsymbol{w}^{(t+1)}\leftarrow r\boldsymbol{w}^{(t)}+(1-r)\boldsymbol{w}^{esk}$$

**For** $k\leftarrow1$ to $K$ **do**

**if** $\boldsymbol{w}^{\left( t+1 \right)}\left( k \right)<\epsilon$ **then**

$$\boldsymbol{w}^{(t+1)}\left( k \right)\leftarrow U\left( \epsilon,\frac{1}{K} \right);$$

**end**

**end**

$\boldsymbol{w}^{(t+1)}\leftarrow\boldsymbol{w}^{\left( t+1 \right)}/\mathrm{sum}(\boldsymbol{w}^{(t+1)})$;

**Supplementary Table 4**. The training of scDEC-Hi-C model. The default settings in all the experiments are as follows. We use Adam optimizer for gradient descent and parameters updating. $\alpha=0.002$ , $m=32$, $n_{d}=5,$the default parameters for Adam optimizer were used. After training scDEC-Hi-C for certain batches (default: 30000), the model will stop training if the cluster proportion reaches a stable distribution.

**Algorithm** Training procedure of scDEC-Hi-C

**Require**: $\theta_{e}^{0}$, $\theta_{d}^{0}$, $\theta_{g}^{0}$, $\theta_{h}^{0}$ for initial parameters of Encoder, Decoder, G and H network, $\theta^{0}=(\theta_{e}^{0}$, $\theta_{d}^{0},\theta_{g}^{0},\theta_{h}^{0})$, $\omega_{dx}^{0}$ and $\omega_{dz}^{0}$for initial parameters of D*_x_* and D_z_ network, $\omega^{0}=(\omega_{dx}^{0},\omega_{dz}^{0})$, batch size $m$ and learning rate $\alpha$.

**While** $\theta, \omega$ have not converged, **do**

**For** $t=1,\ldots,n_{d}$, **do**

**For** $i=1,\ldots,m$, **do**

Sample data $\boldsymbol{x}^{chr} \sim\mathcal{P}_{\boldsymbol{x}^{\boldsymbol{chr}}}$ , latent variable $\boldsymbol{z}\sim\mathcal{P}_{\boldsymbol{z}}, \boldsymbol{c}\sim\mathcal{P}_{\boldsymbol{c}}$ and a random number $\zeta, \eta\sim U(0,1)$.

$\boldsymbol{z}^{chr}\leftarrow$E$\left( \boldsymbol{x}^{chr} \right)$

$\boldsymbol{x}\leftarrow Concat\left( \boldsymbol{z}^{chr} \right)$

$\tilde{\boldsymbol{x}}\leftarrow G\left( \mathbf{z},\mathbf{c} \right)$

$\tilde{\boldsymbol{z}},\tilde{\boldsymbol{c}}\leftarrow H\left( \boldsymbol{x} \right)$

$\hat{\boldsymbol{x}}\leftarrow\zeta\boldsymbol{x+}\mathbf{(}1\mathbf{-}\zeta\mathbf{)}\tilde{\boldsymbol{x}}$

$\bar{\boldsymbol{z}}\leftarrow\eta\boldsymbol{z+}\mathbf{(}1\mathbf{-}\eta\mathbf{)}\tilde{\boldsymbol{z}}$

$L_{ae}^{(i)}\leftarrow{||\boldsymbol{x}^{chr}-D(E(\boldsymbol{x}^{chr}))||}_{2}$

$L_{dx}^{(i)}\leftarrow D_{x}\left( \tilde{\boldsymbol{x}} \right)-D_{x}\left( \boldsymbol{x} \right)+\lambda{({{||\nabla_{\hat{\boldsymbol{x}}}D}_{x}\left( \hat{\boldsymbol{x}} \right)||}_{2}-1)}^{2}$

$L_{dz}^{(i)}\leftarrow D_{z}\left( \tilde{\boldsymbol{z}} \right)-D_{z}\left( \boldsymbol{z} \right)+\lambda{({{||\nabla_{\bar{\boldsymbol{z}}}D}_{z}\left( \bar{\boldsymbol{z}} \right)||}_{2}-1)}^{2}$

$L_{d}^{(i)}=L_{dx}^{(i)}+L_{dz}^{(i)}$

**End**

$\omega\leftarrow\omega+\alpha\cdot Adam(\nabla_{\omega}\frac{1}{m}\sum_{i=1}^{m} L_{d}^{\left( i \right)},\omega)$

**End**

**For** $i=1,\ldots,m$, **do**

Sample data $\boldsymbol{x}\sim\mathcal{P}_{\boldsymbol{x}}$ , latent variable $\boldsymbol{z}\sim\mathcal{P}_{\boldsymbol{z}}, \boldsymbol{c}\sim\mathcal{P}_{\boldsymbol{c}}$.

$\tilde{\boldsymbol{x}}\leftarrow G\left( \mathbf{z},\mathbf{c} \right)$

$\tilde{\boldsymbol{z}},\tilde{\boldsymbol{c}}\leftarrow H\left( \boldsymbol{x} \right)$

$L_{g,h}^{(i)}\leftarrow-D_{x}\left( \boldsymbol{x} \right)-D_{z}\left( \tilde{\boldsymbol{z}} \right)+10{(||\boldsymbol{x}-G(H(\boldsymbol{x}))||}_{2}^{2}+{||\boldsymbol{z}-H(G(z))||}_{2}^{2}+CE(\boldsymbol{c},\tilde{\boldsymbol{c}}))+L_{ae}^{(i)}$

$\theta\leftarrow\theta+\alpha\cdot Adam\left( \nabla_{\theta}\frac{1}{m}\sum_{i=1}^{m} L_{g,h}^{(i)},\theta\right)$

**End** **while**
